# CNVizard – a lightweight streamlit application for an interactive analysis of copy number variants

**DOI:** 10.1101/2024.06.14.598969

**Authors:** Jeremias Krause, Carlos Classen, Daniela Dey, Eva Lausberg, Luise Kessler, Thomas Eggermann, Ingo Kurth, Matthias Begemann, Florian Kraft

## Abstract

Methods to call, analyze and visualize copy number variations (CNVs) from massive parallel sequencing data have been widely adopted in clinical practice and genetic research. To enable a streamlined analysis of CNV data, comprehensive annotation and good visualization are indispensable. The ability to detect single exon CNVs is another important feature for genetic testing. Nonetheless, most available open-source tools come with limitations in at least one of these areas. One drawback is that available tools deliver data in an unstructured and static format which requires subsequent visualization and formatting efforts. Here we present CNVizard, a lightweight streamlit app which requires minimal computational knowledge, and which is compatible with widely used CNV processing tools (CNVkit and AnnotSV). CNVizard can process short- and long-read sequencing data and provides an intuitive webapp-like experience enabling an interactive visualization of CNV data.

## Introduction

Copy Number Variations (CNVs) are a significant aspect of genomic variation, involving amplification or deletion of small or large segments of DNA. These variations contribute substantially to genetic diversity among individuals and populations, and they have been increasingly recognized for their role in the etiology of various genetic diseases.

Historically large CNVs have been mostly analyzed using microarrays, while multiplex ligation-dependent amplification (MLPA) as targeted approach was used for smaller CNVs. In MLPA individual genes are analyzed using a probe mix which is highly specific. While legacy methods such as MLPA and microarray are still used, exome- and genome-wide MPS (massive parallel sequencing)-based analysis of patients with genetic disease became first line diagnostic approach in the recent years and allows comprehensive CNV detection in addition to SNVs(1). Commonly used methods for CNV detection, e.g. microarray or MLPA, are therefore more and more replaced by CNV analysis of MPS data. Accordingly, an increasing number of bioinformatic tools for CNVs analysis in MPS data are available (e.g. CNVkit(2), CNVnator(3), GATK(4)). Most of these tools are suitable for the identification of larger CNVs which compromise several exons of a gene or even larger parts of the genome.

However, in genetic testing and research also single exon alterations must be identified reliably. Some of the available CNV analysis tools also allow the calling of single exon deletions or amplifications, e.g. CNVkit(2). Beside the calling of these variants a comprehensive visualization of the CNV data is important. Nevertheless, most tools lack a comprehensive visualization function, e.g. single exon MLPA/Coffalyzer-like CNV plots, which might be one cause hindering the transition from e.g. MLPA-based to MPS-based CNV analysis. Moreover, a comprehensive annotation of the data with known pathogenetic and database-curated CNVs fosters a fast and reliable analysis. Though several tools are available for CNV calling and/or visualization, most of them lack some of the features, e.g. single exon or family-based analysis, necessary for comprehensive data analysis and visualization in genetic testing and research.

Here we describe CNVizard, a python tool with a browser-based graphical user interface created with streamlit and a Snakemake pipeline (CNVand), to prepare the files necessary for data CNVizard. The CNV analysis is done with the widely used tool CNVkit(2), which allows CNV calling of targeted and genome-wide data down to single exon level. For data annotation we utilized AnnotSV(5).

## Methods and results

CNVizard is written in Python. It provides an interactive browser-based environment, realized as a streamlit application, which enables the structured analysis of CNVs. CNVizard provides filterable data grids using pandas(6) and interactive plots generated by plotly(7).

### A. CNVkit

CNVkit(2) is a python package and command line tool capable of calling CNVs from high-throughput sequencing data down to single exon-level resolution. It belongs to the group of CNV calling algorithms which rely on a read depth-based and B-allele-frequency (BAF) strategy. In brief, these algorithms predict CNVs, by comparing the number of reads of a specific location to the number of reads present in a reference.

### B. AnnotSV

AnnotSV(5) is a tool for annotation of CNV data with additional information which aids the interpretation of pathogenicity.

### C. Data input

#### Mandatory Data input

Depending on the required functionality, CNVizard is designed to work with formatted output provided by CNVkit(2) and AnnotSV(5). The Snakemake(8) workflow provided along with CNVizard, CNVand(16), prepares all necessary files, starting from alignment files. By providing the option to combine an exon-level resolution analysis for CNVs with the flexibility to additionally review annotated VCFs generated by different copy number callers, CNVizard enables an extensive analysis of CNVs.

#### Configurations files

Next to mandatory files, the CNVizard can be modified with configuration files which enable customization of some of its functionalities. These include a tab-delimited text file utilized for formatting the AnnotSV input data, the option to create an env file for an IGV outlink and text files of genes, which enable a panel-based analysis.

### D. Core functionalities

#### Individual Exon-level CNV Analysis

Utilizing the copy number ratio (.cnr) file and the output from the additionally performed bintest provided by CNVkit(2), pandas(6) is used to generate formatted data frames which enable the user to filter seven presets of CNVs on an exon-level. The following presets are available: “total” (represents a formatted version of the unfiltered cnr file), “bintest” (represents a formatted version of the unfiltered bintest output file), “homozygous deletion” (data from the cnr file, filtered for homozygous deletions), “total candidate genes” (data from the cnr file, filtered for CNVs contained inside a candidate gene list), “bintest candidate genes” (data from the bintest output file, filtered with a candidate gene list), “consecutive deletions” and “consecutive amplifications” (data from the cnr file, filtered for consecutively deleted exons; the cut-off value can be set by the user). All dataframes also contain an internal frequency for each predicted CNV, which could be calculated from an internal cohort. The scripts for frequency calculation are provided along with CNVizard. We provide several lists of candidate genes for different genetic conditions; additional ones can be added by the user. For this we provide a script which can be used to transform panel TSV files from the genomics england panel app into compatible TXT files. The user can select the preferred gene-panel list inside the sidebar of the streamlit webapp. Additionally, the user can filter the “total” preset with a variety of filters (“chromosome”,”position”,”gene” etc.). An Overview of the webapplication can be seen in Figure 1.

**Fig. 1.**
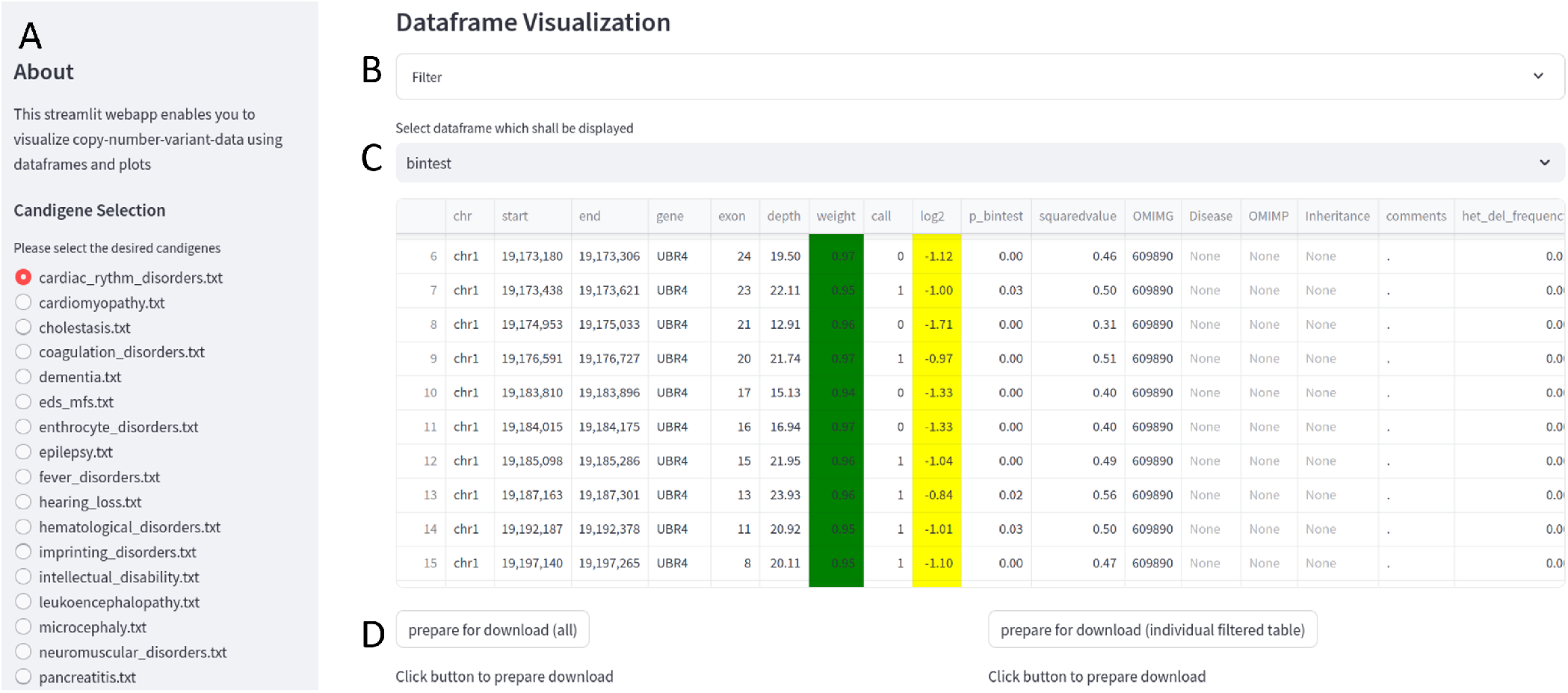
CNVizard web interface. On the sidebar a gene-panel list can be selected. In this screenshot the preset “bintest” has been selected. A deletion of UBR4 is detected (C, column call and log2). A) Sidebar with gene-panel selection; B) Filter-section: drop down menu which enables the user to interactively filter the datagrid; C) Interactive datagrid with color-coding for CNVs (CNV 2 is shown in white, whereas CNV below - 0.65 are marked in yellow); D) Download button, which allows the downloading of the filtered or unfiltered datagrid.

#### Interactive exon-level MLPA-like like log2 and raw depth box-plots

The .cnr file contains two values for the coverage depth. A bias corrected one called “log2 coverage depth” which represents the comparison of region with defined size (bin) coverage depth to the average coverage depth of a pooled reference, with outliers being removed. Additionally, a non-bias corrected coverage depth value called “depth”, which represents the mean coverage depth of the bin. We utilized these two metrics to generate MLPA/Coffalyser-like box plots by computing the median, the mean, upper and low quartile of each exon’s log2 coverage depth and depth value present in the exome or genome reference. With this reference the user can create an exon-level box plot using plotly(7) and plot individual log2 and raw depth values to analyze individual values compare to the reference. These two values are highly dependent, therefore, the box plots should coincide for real CNVs, while false positive calls might only occur in one of the plots. Examples of coverage plots can be seen in Figure 2.

**Fig. 2.**
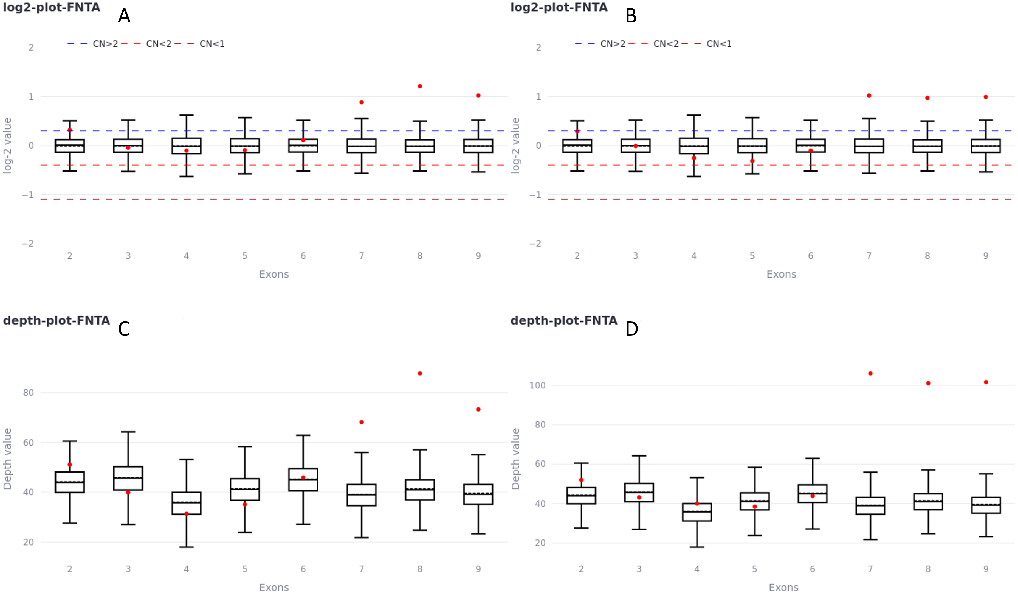
Example MLPA/Coffalyser-like box plot. A deletion of exons 7,8 and 9 of FNTA was observed. Inside the box plots a rise in log2 coverage depth and raw depth was observed (upper panel: The blue and light red dashed line indicate the threshold for a copy number higher or lower than 2. The red dashed line illustrates the threshold for a copy number below 1. Upper panel/lower panel: Box plots indicate the 0.25 and 0.75 quartile. The dashed black lines indicate the mean and the solid black line the median. The whiskers showing the Red dots indicate the copy number or depth of the analyzed sample. A comparison between short read data (A and C) and long read data is shown (B and D))

#### Trio mode

CNVizard also offers an option to provide parental samples which enable a trio analysis, e. g. to filter for *de novo* variants.

#### Genome-wide and chromosome-wide scatter-plot

CNVkit(2) provides functionality to plot the log2 coverage depth values and segmentation calls as a scatter plot. We integrated this functionality into the CNVizard, thereby enabling the user to plot and subsequently analyze CNV data on chromosome or genome level. Examples of scatter plots can be seen in Figure 3.

**Fig. 3.**
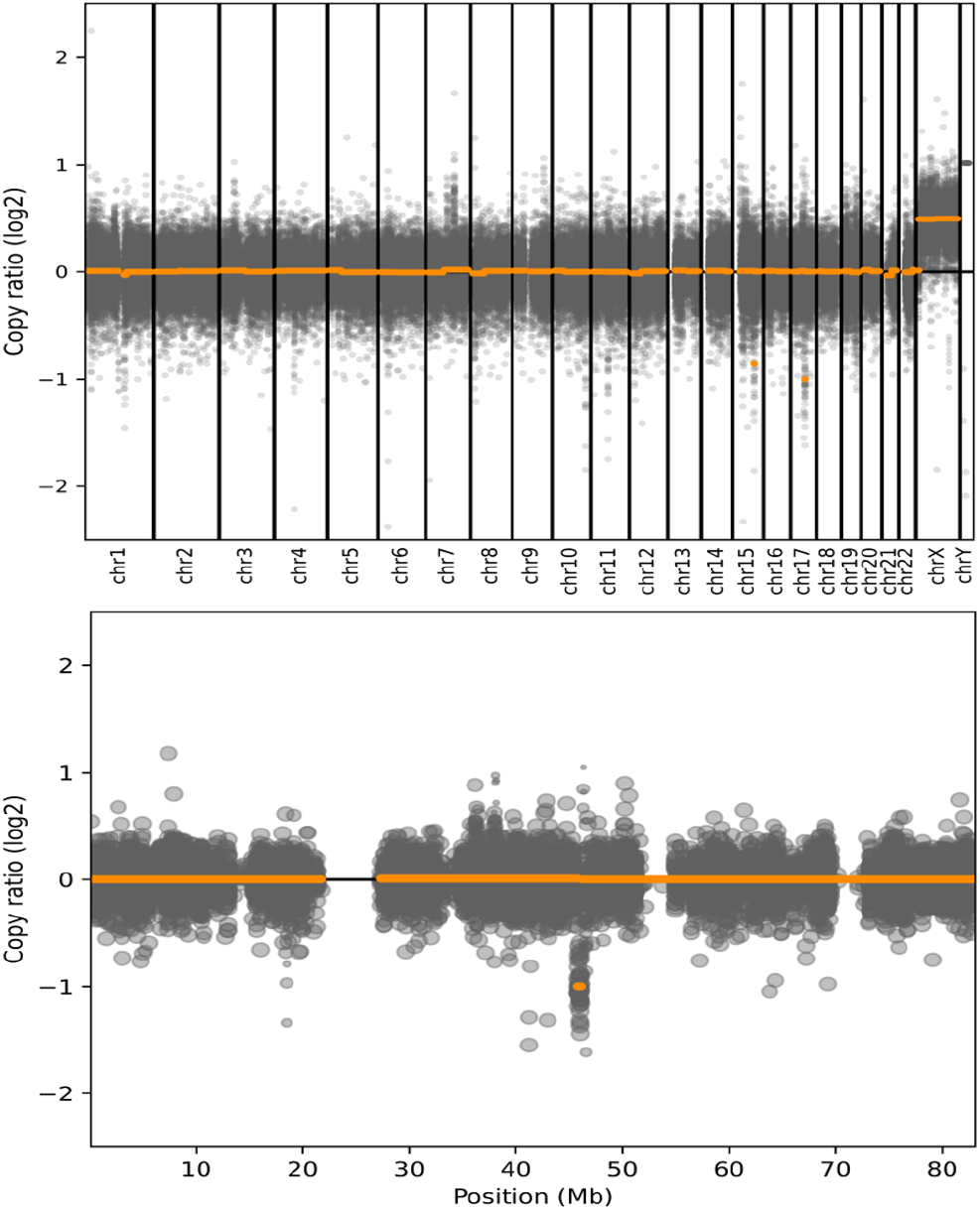
Chromosomal scatter plots as provided by CNVizard. CNVizard uses the scatter plot functionality which is implemented in CNVkit(2) to provide scatter plots of either single chromosomes or the whole genome. CNVkit(2) identified a 564 kb deletion on chromosome 17. The above plot shows the scatter plot of all chromosomes. The zoom-in (lower figure) shows the scatter plot for chromosome 17 where the relevant deletion is easily visible. (Grey dots indicate the copy number of a single bin. Orange dots indicate the copy number of a segment)

**Fig. 4.**
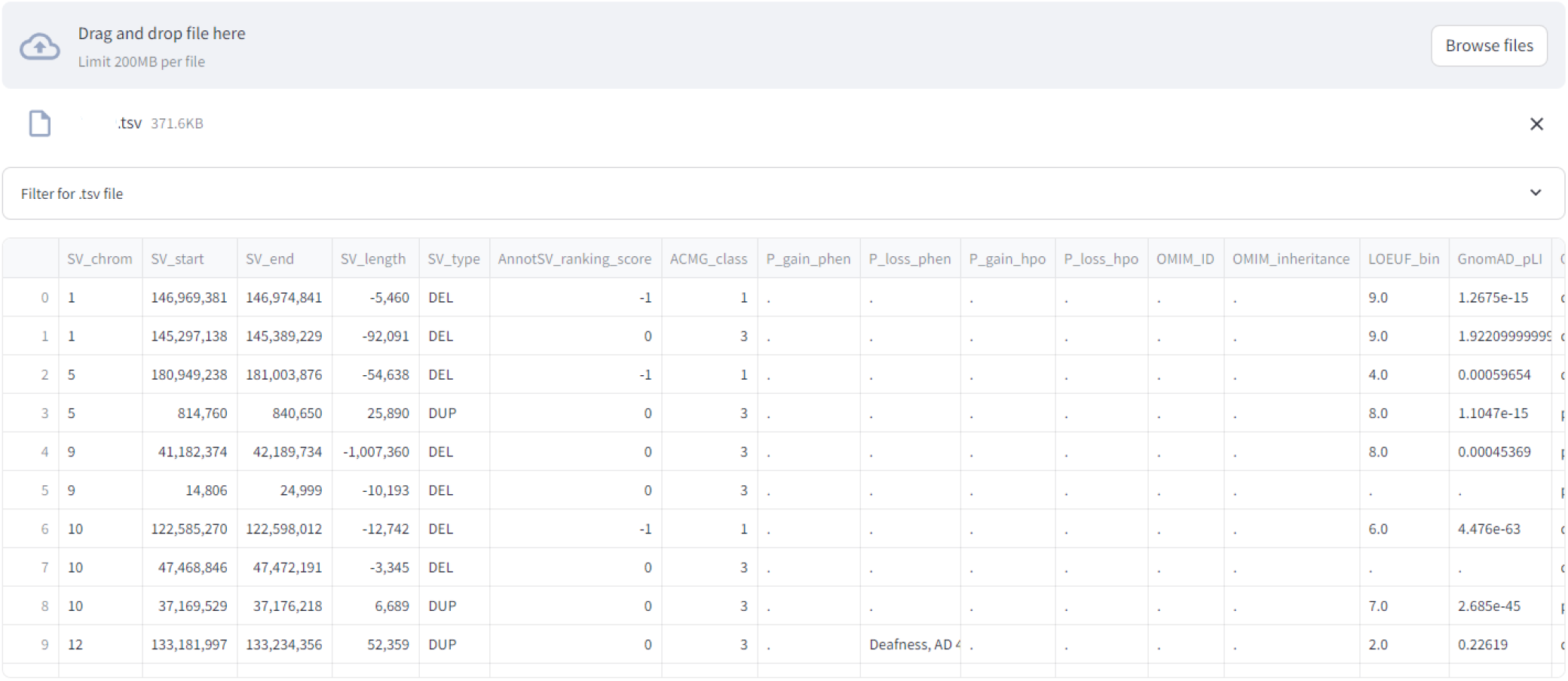
CNV analysis aided by AnnotSV(5): : Depiction of the formatted output provided by AnnotSV.

#### CNV annotation and prioritization

Analyzing CNVs for pathogenicity requires an extensive annotation. Therefore, we integrated support for AnnotSV(5) annotated CNV files, which could be either generated by the CNVand pipeline or other workflows. AnnotSV(5) is compatible with the majority of CNV callers which provide their output in form of a VCF or BED file. Our strategy here was to combine the detection of exon-level CNVs using the CNVkit bintest script and of larger CNVs, called with the CNVkit standard workflow. This functionality complements the more targeted analysis and enables a “discovery mode” for all CNVs in a provided dataset. Next to the required Input files, the user can provide a text file which limits the very comprehensive AnnotSV(5) output file, to a smaller subset of annotations.

#### Snakemake-Pipeline

Next to the CNVizard we provide a Snakemake pipeline(8) called “CNVand” which implements the preprocessing with CNVkit(2) and the annotation with AnnotSV(5), starting from alignment and variant call files. CNVand is compatible with panel, exome and genome sequencing data. The Snakemake(8) pipeline is available through the Snakemake Workflow Catalog(8) and as a package on GitHub.

#### Generating internal references

To obtain frequencies from internal cohort data we wconcatenated .cnr files using pandas(6) and subsequently calculated exon-level frequencies. We provide frequencies of our exome and genome cohorts as reference files and the scripts needed to create such internal frequencies on own data.

## Discussion

Whereas tools for CNV calling and visualization have been developed and published previously, to our knowledge none of them combines the possibility to analyze CNVs ranging from single exon up to whole genome in a streamlined process. To pinpoint differences and advantages of CNVizard compared to already available open-source tools, we addressed different aspects which are important for a streamlined CNV analysis. The first criterion is the ability to robustly call CNVs, therefore enabling a workflow independent of an existing pipeline. Along with 3 out of 5 other tools the CNVizard can perform independent CNV calling with the CNVand Snakemake(8) pipeline. Whereas CNspector(9) and iCopyDAV(10) implement their own CNV calling algorithms, CNVizard utilizes the widely used and actively maintained tool CNVkit(2) for CNV calling. A second important criterion is the visualization of the CNV data and therefore the integration of an interactive datagrid. While all tools implemented a datagrid in some form, only knotAnnotSV(11) and CNVizard provide filter options to individually prepare the data for the analysis. Both tools provide flexible filtering options and contain sufficient annotations presented in a structured format. While most tools (4 out of 5) have the option to analyze the ingested data for larger structural alterations using a scatter plot, only CNVizard enables plotting for smaller/exon-level alterations. Additionally, the majority of previously published tools (4 out of 5) respect only sparse possibilities for annotation, some of them come with integrated annotation sources, which are susceptible to be outdated, if not properly maintained. By providing support for AnnotSV(5), which is a widely used and actively maintained framework for the annotation of CNVs, CNVizard can provide an exhaustive number of annotations for larger CNVs and supports a more condensed number of annotations (inhouse frequency, OMIM-annotations and Inheritance) for smaller CNVs. The importance of up-to-date resources for annotation have been already demonstrated(12). Depending on the type of genetic testing or research focus, only a few genes could be important for the analysis. Gene panels are only implemented by the minority of previously published tools or require a reformatting of the input data. To overcome this limitation, CNVizard has a straightforward implementation of gene panels, which could be easily adjusted by the user. Furthermore, we compared the tools for their capability of performing an analysis for loss-of-heterozygosity. We did not implement this feature in the CNVizard, since it is mostly used in the analysis of somatic CNVs. Nevertheless, 2 out of 5 tools provide this feature, and it might be a useful addition to the CNVizard for larger CNVs and for uniparental disomies (UPD). One of the important features of CNVizard is the ability to analyze single exon CNVs. To our knowledge no other tool provides such high resolution in an interactive datagrid. Furthermore, CNVizard provides, similarly to MLPA analysis, box plots for log2 transformed CNV and sequencing depth for single exons (Figure 2). Genetic testing and research are often done in family-based setup, e.g. as trio analysis of the affected individual with respect to parental data. The ability to analyze such cases together in a family mode is a crucial feature, which is supported by CNVizard and only 2 out of 5 comparable tools. Basically, all tools and CNVizard are also compatible with other CNV callers. By relying on AnnotSV(5) as an annotation tool, which is compatible with a variety of different CNV-calling algorithms, CNVizard inherits this compatibility in the context of VCF files. CNVizard is easy to set up, as it is open source and available via GitHub. The tool is provided as a python package, which installs all dependencies automatically. Its unique feature is the comprehensive implementation of a CNV analysis environment which offers a high-resolution analysis of CNVs, which is a relevant topic in the research of monogenetic diseases. CNVizard offers a pipeline for CNV calling (CNVand), starting from alignment files, offers an interactive datagrid with various filter options, allows the analysis and visualization of single exon CNV, similar to MLPA/Coffalyser analysis, a comprehensive configurable annotation, gene panel-based filter strategies, trio analysis and is compatible with other CNV callers, beside CNVkit.(2).

In summary, CNVizard is a lightweight CNV analysis toolkit which enables a comprehensive analysis of CNV data for diagnostic and research applications.

## AUTHOR CONTRIBUTIONS

JK, IK, MB and FK conceived the Idea. JK, CC, DD, MB and FK implemented the streamlit application and the python analysis. CC and FK implemented the snakemake workflow. EL, LK and TE performed user testing. JK, CC, MB, TE and FK performed validation. JK and FK wrote the manuscript. IK, TE, MB and FK supervised the project. All authors reviewed the manuscript and gave their consent for the submission of the final version.

## CODE AVAILABILITY

All code to setup CNVand (https://github.com/IHGGM-Aachen/CNVand) and CNVizard (https://github.com/IHGGM-Aachen/CNVizard) is available on Github. Additionally CNVand is available on WorkflowHub(16).

## ACKNOWLEDGEMENTS

This research project was funded by the START-Program of the Faculty of Medicine. We thank Ricardo Henriques and his group for the distribution of their biorxiv latex template via overleaf https://de.overleaf.com/latex/templates/henriqueslab-biorxiv-template/nyprsybwffws.

---

**Table 1.** Comparison between CNVizard and other CNV analysis tools.

| Functionality provided | CNSpector(9) | reconCNV(13) | CNViz(14) | GenomeCAT(15) | knotAnnotSV(11) | CNVizard |
| --- | --- | --- | --- | --- | --- | --- |
| CNV-Calling | Yes | Yes | No | Yes | No | Yes |
| Interactive Datagrid | Yes | Limited | Limited | Limited | Extensive | Extensive |
| Scatterplot (genomewide) | Yes | Yes | Yes | Yes | No | Yes |
| Boxplot (gene/exon-based) | No | No | No | No | No | Yes |
| Annotation | Depends on Input Data | limited | limited | limited | Extensive | Extensive |
| Panel based Analysis | Yes | No | No | No | No | Yes |
| Loss of Heterozygosity | Yes | No | Depends on Input Data | No | No | No |
| Single-Exon Resolution | Limited | No | No | No | Depends on Input Data | Extensive |
| Family / trio mode | Yes | No | No | Yes | No | Yes |
| Compatibility with third party CNV-Callers | Yes | Yes | Yes | Yes | Yes | Yes |
| Architecture | R/Shiny App | Python/env | R/Shiny App | Java / Installation Wizard | Perl | Python/env/streamlit |

